# NMDAR-mediated shift of neuronal gain across the cortical hierarchy

**DOI:** 10.64898/2026.03.04.709528

**Authors:** Movitz Lenninger, Pawel Herman, Mikael Skoglund, Arvind Kumar

**Author notes:** Contact: Arvind Kumar.

## Abstract

Neuronal gain is a critical feature of neuronal information processing, reflecting a cell’s excitability in response to presynaptic inputs. Evidence from spine counts in non-human primates suggests that the number of presynaptic inputs increases across the cortical hierarchy. Therefore, understanding the regulation of the neuronal gain can yield important insights into different integration strategies across the cortex. Along with the increasing spine counts, there is further evidence for a similar increase in NMDA-receptor-mediated currents. Here, we show, using detailed simulations of a thick-tufted layer 5 pyramidal neuron, that the amount of current mediated through NMDA receptors (NMDARs) strongly affects the neuronal gain function with balanced presynaptic inputs. Without NMDA, postsynaptic activity quickly saturates and declines with increasing inputs. However, with NMDARs, the gain of postsynaptic activity shifts into higher input regimes, enabling integration of dense inputs. We show that, with NMDARs, a significant fraction of postsynaptic activity arises from feedback between the voltage dependence of NMDARs and voltage-gated sodium channels. Lastly, we show, using a network model, that NMDAR knockout in single cells has differential effects on postsynaptic activity depending on network size (a proxy for cortical hierarchy). Thus, we propose that an NMDA gradient across the cortical hierarchy may be necessary to maintain excitability amid increasing presynaptic input.

## Introduction

The neuronal gain is a critical feature controlling network dynamics and the integration of presynaptic inputs. In-vivo recordings suggest that, in active network states, neurons receive large amounts of excitatory and inhibitory synaptic inputs across the dendritic tree, causing strong depolarizations and increased variability with respect to the resting state (1). Both experimental and computational studies have shown that synaptic background inputs can modulate the gain of a neuron (2; 3; 4; 5). Prior work has also suggested that when the excitatory and inhibitory inputs are balanced, high synaptic input rates can paradoxically lead to a reduction in postsynaptic activity (6). Importantly, across the cortical hierarchy of primates, there is a trend of increasing spine counts along the basal dendrites (7; 8; 9; 10). In the macaque prefrontal cortex, the number of basal spines in the prefrontal cortex (PFC) is approximately 10 times larger than in the primary visual cortex (V1). Because the number of spines is a proxy of synaptic input, this suggests that many higher cortical areas might operate in high input regimes. Therefore, given the paradoxical decrease in postsynaptic activity at high input states, it is important to identify and understand mechanisms that may adapt the neuronal gain function to different input regimes.

AMPA (*α*-amino-3-hydroxy-5-methyl-4-isoxazolepropionic acid) and NMDA (N-methyl-D-aspartate) receptors (AM-PAR and NMDAR, respectively) are two major sources of excitatory currents in the brain, but they strongly differ in kinetics, voltage-dependence, and integrative properties. AMPAR-mediated conductances rapidly rise and decay upon presynaptic inputs, while NMDAR-mediated conductances have considerably slower kinetics and scale with the local membrane potential due to Mg^2+^-block (11; 12; 13). The NMDA/AMPA ratio varies across age, cell type, and brain area (14; 15, and see Table 1 in (16)). The NMDA/AMPA ratio can also be modified by activity-dependent calcium-based plasticity, which primarily up-or down-regulates AMPARs (17). Furthermore, evidence based on gene expression in primates suggests an upregulation of the NMDA subunit GluN2B across the cortical hierarchy (18; 10). As subunit GluN2B is associated with an increased NMDAR decay time compared to subunit GluN2A (19; 20), this upregulation suggests a coinciding increase of NMDAR-mediated current in higher cortical areas. Similarly, studies report higher ratios of GluN2B/GluN2A and longer NMDAR decay time constants in rat PFC compared to V1 (21), indicating that the relative amount of NMDAR-mediated current is higher in the rat PFC despite the peak NMDA/AMPA ratio being similar across the two regions (16).

Apart from being crucial for calcium-based long-term plasticity (22; 17), NMDARs have also been linked to NMDA spikes (or plateau potentials) (23; 24), high-frequency bursting (25), persistent activity (26), increased action potential generation during evoked activity (27), increased orientation selectivity (28), improved sequence discrimination (29), differential integration strategies across inhibitory cell-types (15), etc. Given the voltage-dependence of NMDARs, we hypothesize that an increase in NMDAR-mediated current can also shift neuronal gain into a high-input regime.

To test this hypothesis, we first simulate responses in a detailed biophysical model of a thick-tufted layer 5 (TTL5) pyramidal neuron. Indeed, we find that the inclusion of NMDARs reshapes the neuron’s input-output transfer function. In particular, NMDARs allow the neuron to generate strong postsynaptic activity in response to strong presynaptic input. However, for sparse inputs, including NMDARs instead reduces postsynaptic activity. Thus, varying the amount of NMDAR-mediated current makes the neuronal gain sensitive to different levels of presynaptic input. Finally, we show, in network simulations (using single-compartment neuron models), that single-cell NMDAR knockout is expected to increase postsynaptic firing rates in small networks (proxy for low cortical areas) but decrease them in large networks (proxy for high cortical areas). Based on these results, we argue that the increase in NMDAR-mediated currents across the cortical hierarchy is an important mechanism allowing neurons to integrate inputs across an increasing number of synapses.

## Results

To explore the role of NMDARs on the spiking activity of neurons, we consider the responses of a single detailed reconstruction of a TTL5 pyramidal cell (30) with excitatory and inhibitory synapses (Fig. S1A, see Methods for details). All excitatory synapses consist of a mix of AMPARs and NMDARs, which are assumed to respond to the same presynaptic inputs (31; 32). The NMDA/AMPA ratio, *α*_*N/A*_, is defined as the ratio of peak conductances in the absence of Mg^2+^-block (Eq. 3; Fig. S1B). Unless otherwise specified, we assume the NMDAR conductance decay time to be 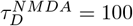 ms and the NMDA/AMPA ratio to be 1.1 (w/ NMDA) (21) or 0 (w/o NMDA).

A striking property separating NMDARs from AMPARs is the voltage-dependence of NMDARs (13), implying that the effective synaptic strengths of NMDARs and AMPARs are differently modulated by the local membrane potential where the receptors are situated (Fig. S1C). Increasing the synaptic inputs leads to increased depolarizations of the membrane potentials across the cell (Fig. S1D-E). Thus, the effective synaptic strength of NMDARs dynamically increases with increasing input levels and distance from the soma. Therefore, for each choice of NMDA/AMPA ratio (*α*_*N/A*_) or NMDAR decay time 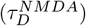, we uniformly scaled the synaptic weights such that the unitary excitatory charge is equal at *V*_0_ = −54.5 mV (Fig. S1C, dashed line). *V*_0_ was chosen such that the mean subthreshold, measured without voltage-gated sodium (Na^+^) channels, somatic membrane potentials, and the standard deviations are comparable across *α*_*N/A*_ = 0 and *α*_*N/A*_ = 1.1 (Fig. S1F-G). The synaptic strengths (*Q*_0_) are set such that the average peak somatic EPSC is ∼17 pA from basal synapses (w/ NMDA) when measured at −70 mV, consistent with data reported by (21) (see Methods for details). Unless otherwise stated, we assume that the total, time-averaged inhibitory conductance across the cell is twice the excitatory conductance, E/I ratio = 0.5, which gives comparable and in-vivo-like subthreshold potentials (Fig. S1F-G). Note that, in the following, synaptic input rates are only reported by the total excitatory input rates across the entire cell, but there is a corresponding increase in inhibitory inputs, maintaining the E/I ratio.

### Input-dependent excitability is modulated by NMDARs

To study how the dynamic regulation of effective synaptic weights via NMDARs influences neuronal excitability, we first measure somatic input resistance across several input conditions. First, we measure the input resistance in the absence of Na^+^ channel conductances to avoid interference from action potentials. To reduce measurement noise due to fluctuating synaptic inputs, we first record somatic potentials without current pulses; then, using the same synaptic inputs, we measure the potential while applying current pulses to the soma (40 pA, roughly twice the size of a single EPSC). Lastly, we calculate the difference, Δ*V*, between the two conditions (Fig. 1A). This allows us to reliably estimate the input resistance for each current pulse, *R*_*in*_ = Δ*V/I* (*I* is the applied current). Even without synaptic inputs, the input resistance at the soma is low (∼ 40 MΩ) (Fig. 1B, *λ*_*E*_ = 0 kHz). Lower somatic input resistances (∼ 20 MΩ) have been reported for layer 5 pyramidal cells in the somatosensory cortex of adult (P56) rats (33), while larger input resistances (∼ 70 MΩ) have been reported in rat PFC (P30) (34). The input resistance decreases further with increasing synaptic inputs, reaching in-vivo-range values (35) (Fig. 1B). However, the measured input resistance is consistently larger with NMDARs (*α*_*N/A*_ = 1.1) than without NMDARs (∼ 75% larger at high-input conditions). Furthermore, tracking the spatial depolarization across each cell reveals that including NMDARs gives (1) stronger depolarizations across the cell, and (2) longer characteristic length scales (Fig. 1C). The dendritic depolarization is primarily confined to the basal dendrites (Fig. 1C, inset).

**Figure 1.**
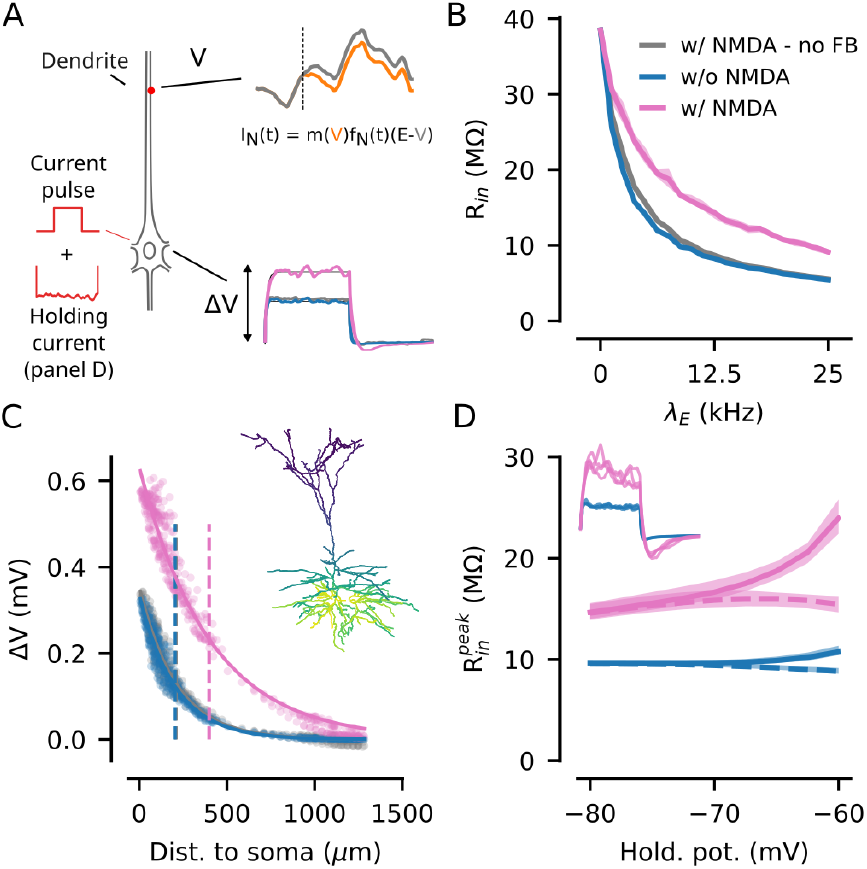
Input-dependent excitability is modulated by NMDARs. **A**) Top: Calculation of NMDAR-mediated current without feedback at an excitatory synapse. Voltage traces with (grey) and without (orange) somatic current pulse (dashed line: pulse onset). The equation governing the NMDA-mediated current is shown, with the respective voltage terms color-coded. Bottom: Examples of somatic depolarizations (Δ*V*) to injected current pulses. **B**) Estimated input resistances versus synaptic input rates (without Na^+^ channels). **C**) Compartment depolarizations due to a single current pulse (*λ*_*E*_ = 12.5 kHz). Solid lines show exponential fits, and dashed lines show the characteristic lengths. Inset: Δ*V* visualized across the cell (*α*_*N/A*_ = 1.1). **D**) Estimated input resistance at different holding potentials with (solid lines) and without (dashed lines) Na^+^ channels (*λ*_*E*_ = 12.5 kHz). Inset: depolarizations across trials (holding potential −60 mV). Panels A) and D): solid lines indicate the mean and shaded areas the min/max values (across 4 trials). The injected current amplitudes are 40 pA (panel B) and 10 pA (panel D) for a duration of 100 ms.

### Increased NMDAR response leads to higher input resistance

Measuring input resistance using current injections relies on the assumption that the injected current does not alter the conductance state of the neuron. However, because NMDARs exhibit voltage-dependent gating, this assumption no longer holds in the presence of presynaptic inputs (e.g., see Supp. Eq. S10 for approximation with NMDARs in a single-compartment model). Thus, the increased input resistance with NMDARs (Fig. 1B) may arise either from differences in effective synaptic weights before (i.e., different background states) or in response to the injected current. To rule out that the differences are not due to different effective synaptic weights before the current injections, we remove the feedback between the injected somatic current pulse and the voltage-dependence of the NMDARs (Eq. 2). This is done by first recording the voltage potentials at each compartment during the trials without the somatic current pulses (Fig. 1A, top, orange line) and then using these recordings to determine the voltage-dependent scalings, *m*(·),of the NMDARs in the trials with the currents applied (Fig. 1A, top, grey line). This has a striking effect, the input resistance (Fig. 1B, gray) and the characteristic length (Fig. 1C, gray) are now close to the values obtained without NMDARs. Thus, removing the feedback between NMDARs and the current injection reveals that there is no large difference in the effective synaptic weights before the currents are injected but, although the somatic depolarizations are small (Fig. 1C), the current injections slightly strengthens the conductances of the NMDARs leading to a stronger depolarizations and therefore larger measured input resistances (Fig. 1B, pink). An analysis of input resistance in single-compartment models (Supp. Eq. S10) suggests an identical change in input resistance with NMDARs in response to small hyperpolarizing currents, which we validated in the TTL5 model (Fig. S2).

### Strong voltage-dependence with NMDARs and Na^**+**^channels

Experimental recordings have shown that the input resistance of single cells is voltage-dependent, which has been attributed to various intrinsic voltage-gated ion channels (36; 37). Next, we show that a similar voltage-dependent feedback between NMDARs and Na^+^ channels can further increase the input resistance at depolarized states. In particular, we investigate the effect of different holding potentials on input resistance in the presence of Na^+^ channels and synaptic inputs. To eliminate the risk of spontaneous action potentials affecting the results, we first record the required somatic “holding current” needed to keep the somatic potential fixed at the desired potential using a voltage clamp (Fig. 1A, holding current). Then, again using the same presynaptic inputs, we replace the voltage-clamp with a time-varying current-clamp, applying the recorded holding current, and track somatic depolarizations during the somatic current pulses. The inclusion of Na^+^ channels across the cell introduces stronger (but still subthreshold) transient depolarizations, especially with NMDARs (Fig. 1D, inset). To capture these transients, as they can be important for generating action potentials, we calculate the peak input resistance, 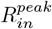, using the maximal somatic depolarization from each current pulse. A small voltage-dependent effect is present both with and without NMDARs, but is significantly stronger when including NMDARs (Fig. 1D, compare blue and pink solid lines). Because the effect is much smaller without NMDARs, the Na^+^ channels alone cannot account for the large changes in input resistance observed with NMDARs. Yet, when repeating the experiment without Na^+^ channel conductance, the effect also largely disappears both with and without NMDARs (Fig. 1D, dashed lines). Combining these observations shows a strong positive feedback between Na^+^ channels and NMDARs. Furthermore, in the presence of Na^+^ channels, the gain in input resistance with NMDARs depends not only on the presynaptic inputs but also on the depolarization of the cell.

### NMDA receptors reshape the input-output function

Next, we study the effect of NMDARs on the input-output transfer function during synaptically evoked activity. As before, unless otherwise stated, we maintain a constant E/I ratio of 0.5 across all input conditions. Without NMDARs, the cell’s firing rate initially increases rapidly with increasing synaptic inputs but eventually starts to decrease (Fig. 2A, blue opaque line). With synaptic NMDARs, however, the cell is insensitive to low synaptic input rates and only becomes active at higher rates (Fig. 2A, pink opaque line). The strong initial gain without NMDARs is likely due to the rapid depolarization of the cell, which is significantly slower with NMDARs (Fig. S1F or Fig. S3A).

**Figure 2.**
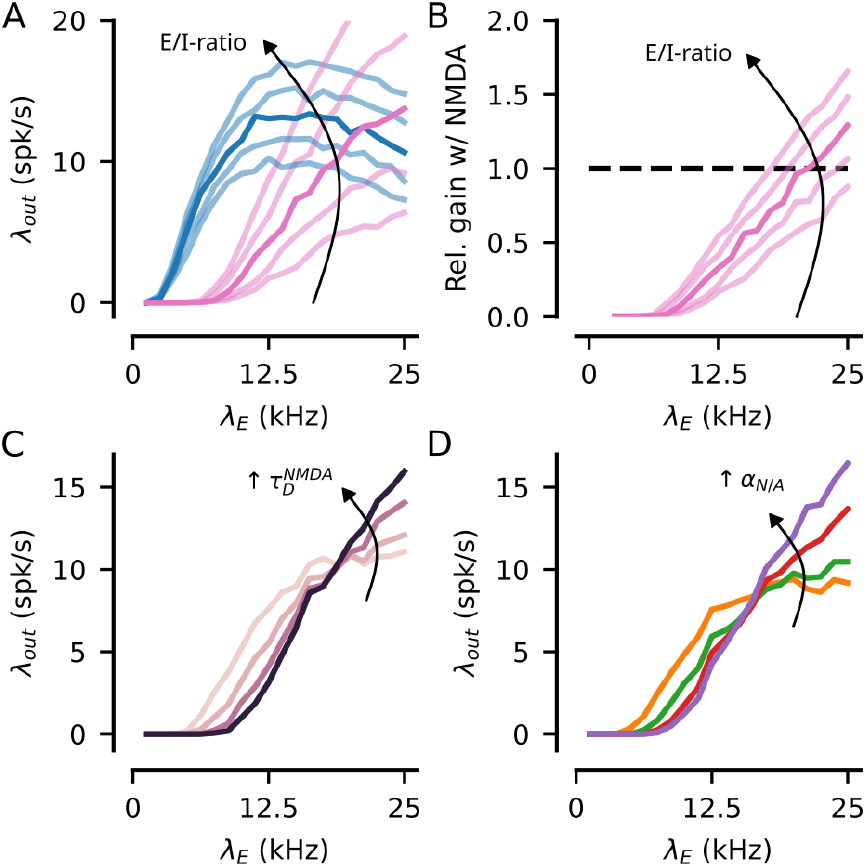
NMDA receptors reshape the input-output function. **A**) Postsynaptic activity as a function of total excitatory input rate (blue - w/o NMDARs, pink - w/ NMDARs). Opaque lines: I/E ratio = 2; transparent lines: I/E ratio ∈ *{*1.9, 1.95, 2.05, 2.1*}* (arrows indicate increasing E/I ratios). **B**) Relative gain of including NMDARs. A relative gain of 1 indicates equal postsynaptic rates with and without NMDARs. **C**) postsynaptic activity for different choices of NMDAR conductance decay times, 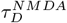 ∈ *{*25, 50, 100, 200*}* ms (*α*_*N/A*_ = 1.1). **D**) postsynaptic activity for different choices of peak NMDA/AMPA ratios, *α*_*N/A*_, ∈ *{*0.25, 0.5, 1, 2*}* (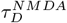 = 100 ms). Each data point is based on 60 simulated seconds.

In vivo measurements typically report that inhibitory conductances are 2-6 fold larger than excitatory (35; 38). Because the effective synaptic weights scale differently with voltage with and without NMDARs, changes in the excitatory/inhibitory ratio (E/I ratio) should differently affect the postsynaptic firing rates. Comparing the output firing rate across different E/I ratios, defined as the ratio of the total excitatory to inhibitory conductances, shows that the cell with NMDARs is indeed more sensitive to the E/I ratio (Fig. 2A, compare blue and pink traces). With NMDARs, the cell depolarizes strongly with increasing input rates for high E/I ratios (Fig. S3A-B), leading to high postsynaptic rates (Fig. 2A, pink traces). On the other hand, at low E/I ratios, reduced depolarizations result in smaller postsynaptic responses. Without NMDARs, however, the influence of the E/I ratio is comparably small. Thus, the relative gain of including NMDARs strongly depends on the E/I ratio (Fig. 2B).

Similarly, varying the NMDARs decay time constant (Fig. 2C), or the peak NMDA/AMPA ratio (Fig. 2D), while preserving the integrated charge at *V*_0_, shows that the change in input-output transfer curve is gradual with respect to the relative amount of NMDAR-mediated conductance (see Fig. S3C-F for subthreshold statistics). Thus, longer NMDAR decay times or larger NMDA/AMPA ratios promote activity during high network activity states but suppress activity during low network activity states.

### NMDAR - Na^**+**^channel feedback increases postsynaptic activity

Next, we ask what factors influence the decrease in postsynaptic firing rates without NMDARs and the strong responses with NMDARs at high synaptic input rates? To check if the differences in subthreshold activity at higher input rates (Fig. S1F-G and Fig. S4A-B) are sufficient to explain the difference in postsynaptic activity, we construct a neuron model where the feedback between the voltage-dependence of NMDARs and the Na^+^ channels is removed. To do so, we first record the subthreshold activity of all compartments during synaptic bombardment without any Na^+^ channel conductances. Then, we reuse these voltage-traces to control the voltage-dependent terms of the NMDAR conductances, Eq. 2, in the presence of Na^+^ channels. Thus, the NMDARs influence the Na^+^ channel conductances, but not vice versa (Fig. 3A). Interestingly, without feedback between NMDARs and Na^+^ channels, the postsynaptic firing rates are consistently lower than without NMDARs (Fig. 3B, compared blue and gray traces). Thus, the difference in subthreshold variability with and without NMDARs alone is not a sufficient explanation of the high postsynaptic firing rates with NMDARs. Instead, NMDAR - Na^+^ channel feedback is crucial for high postsynaptic activity at high input rates, and increases activity by more than 60% relative to without feedback (Fig. S4C).

**Figure 3.**
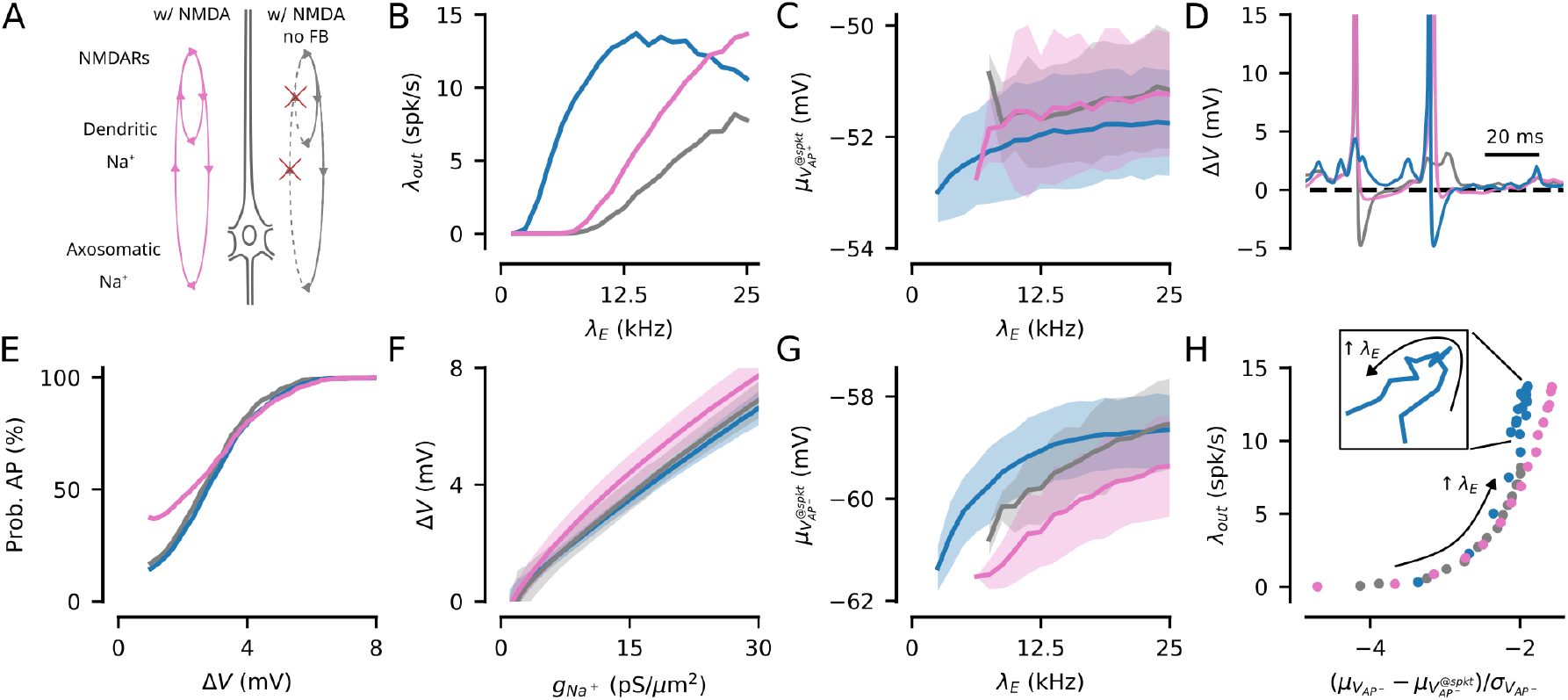
NMDAR - Na^+^ channel feedback increases postsynaptic activity. **A**) Illustration of the difference between ‘w/NMDA’ and ‘w/NMDA no feedback’ conditions. **B**) The input-output transfer function w/o NMDARs (blue), w/ NMDARs (pink), and w/ NMDARs no feedback (gray). **C**) Estimated spiking thresholds in AP^+^ conditions. **D**) Voltage depolarization in the AIS due to Na^+^ channels. **E**) Probability of eliciting an action potential after crossing a certain depolarization, Δ*V*. **F**) Voltage depolarization as a function of Na^+^ channel conductance in the AIS. **G**) Estimated required depolarizations (in the AIS) from synaptic inputs to elicit action potentials. **H**) The relative distance to the synaptically driven threshold provides qualitative information about postsynaptic firing rates. Arrow indicate direction of increasing presynaptic inputs. All panels are based on 60 seconds of simulated time. Panels C), F), and G): Solid lines show mean values, and shaded areas indicate *±* 1 standard deviation.

### Smaller synaptic depolarizations required for AP with NMDA

As the differences in somatic subthreshold potentials are not sufficient to explain the differences in postsynaptic firing rates, we seek the mechanism by which the NMDAR - Na^+^ channel feedback boosts activity. First, we quantify the action potential threshold in the axon initial segment (AIS), the primary site of action potential generation (39; 40) (Fig. 3C). Despite higher input resistance, the estimated thresholds are slightly higher with NMDARs (average spiking thresholds: −52.1 mV w/o NMDARs, −51.5 mV w/ NMDARs, and −51.4 w/ NMDARs but w/o feedback). The lower threshold without NMDARs can be linked to rapid depolarizations driven by AMPAR kinetics, lowering the spiking threshold (41). Thus, differences in observed spiking thresholds do not explain the high firing rates observed with NMDARs at high input rates.

However, the observed spiking thresholds include the depolarizing effect of partial Na^+^ channel openings prior to the estimated spike onset. Therefore, we next aim to disentangle the depolarizations caused by synaptic inputs and by Na^+^ channels. To separate these, we simulate the subthreshold responses (without Na^+^ channels, referred to as AP^−^ conditions) to identical presynaptic spike trains as used in Fig. 3B (which in the following is referred to as AP^+^ conditions). Comparing membrane potentials across these conditions allows us to link action potentials to depolarization caused by synaptic activity and to Na^+^ channels, respectively.

First, by comparing the traces with (AP^+^) and without (AP^−^) Na^+^ channels (Fig. S4D, solid and dashed lines respectively), we can obtain the depolarization, Δ*V*, caused by the opening of Na^+^ channels while largely canceling out the wave-forms caused by presynaptic inputs. Interestingly, we observe that some depolarizations lead to action potentials with NMDARs but not without feedback or without NMDARs (e.g., Fig. 3D). To systematize this observa-tion, we compute the probability of eliciting an action potential after crossing (and before returning to) a particular depolarization, Δ*V*. Indeed, with NMDARs, there is a higher probability of eliciting an action potential after small depolarizations, Δ*V* (Fig. 3E, pink). No large difference is observed without NMDARs and with NMDARs but without feedback (Fig. 3E, blue and gray).

To explain the difference in AP probability, we compare the Na^+^ channel conductance in the AIS against the corresponding Na^+^-induced depolarizations, Δ*V* (Fig. 3F). This reveals stronger depolarizations with NMDARs (Fig. 3F, pink), as predicted by the increased input resistance (Fig. 1B and D). As a control, comparing the Na^+^ channel conductance to the membrane potentials in the AP^+^ condition reveals little difference between the synaptic models (Fig. S4E). Thus, with NMDAR-Na^+^ channel feedback, small depolarizations caused by presynaptic inputs cause increased depolarization due to the partial opening of Na^+^ channels.

Next, we investigate whether the presynaptic depolarization required for APs differs across the cells. We define the “synaptically driven spiking thresholds”, 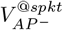, as the membrane potentials in the AP^−^ conditions (i.e., without Na^+^ channels) at the times of spiking in the AP^+^ conditions (i.e., with Na^+^ channels). Surprisingly, despite the higher spiking thresholds in direct measurements (Fig. 3C), the estimated “synaptically driven thresholds” reveal that smaller synaptic depolarizations are needed to generate APs with NMDARs (Fig. 3G). This discrepancy is explained by the additional contribution of partial openings of Na^+^ channels before a “full” AP is generated. This holds both with and without NMDAR-Na^+^ channel feedback (except for at high input rates without feedback), but is consistently stronger with feedback. Thus, while the spiking threshold appears higher in direct measurements with NMDARs, the required synaptically driven depolarization is actually lower, which explains the higher activity with NMDARs at high input rates.

#### A simple heuristic predicts decreasing activity without NMDARs

Unfortunately, it is very difficult to obtain good mathematical descriptions of the firing rate given conductance-based synapses, especially with long conductance decays and with non-linear voltage-dependencies. However, given our estimates of the required synaptically driven depolarizations for action potentials, 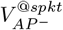, we can compute the distance between the mean membrane potential without Na^+^ channels, 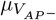, and the mean synaptically driven threshold, 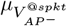, relative to the size of the subthreshold fluctuations, 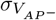 (Fig. S4F-G). Heuristically, the smaller this relative distance is, the larger the postsynaptic activity should be. Although it’s a heuristic predictor, it provides qualitative information about the spiking activity of the cells (Fig. 3H). Without NMDARs, the observed decrease in postsynaptic activity occurs when the difference in mean membrane potential and the required synaptically driven threshold saturates (Fig. S4F, blue), causing the decrease in variability to increase the relative distance to threshold (Fig. S4G, blue). With NMDARs, on the other hand, the slow increase in mean membrane potential implies that the relative difference to threshold decreases despite the reduction in variability, causing a continuous increase in activity across the range of tested inputs. This holds both with and without feedback. However, the lower threshold with NMDAR-Na^+^ channel feedback results in consistently shorter relative distances and, consequently, larger postsynaptic activity.

### The neuronal gain predicts differential effects of NMDAR knockout across the cortical hierarchy

Our single-cell analysis suggests that neurons lacking NMDARs should lose excitability when embedded in large networks with many synaptic inputs, due to reduced postsynaptic activity during high-input conditions. Consequently, the effect of NMDAR knockout should vary across the cortical hierarchy, as the number of recurrent synapses increases. To study the impact of NMDAR knockout across the cortical hierarchy, we model the impact of network size on the population activity in sparsely connected networks of single-compartment neurons (Fig. 4A). Given a fixed connection probability, larger networks imply more recurrent connections, therefore smaller (larger) networks can be considered equivalent to early sensory regions (higher cognitive regions). To simplify the simulations, we assume an equal number of identical excitatory and inhibitory neurons, that all recurrent excitatory synapses have an NMDA/AMPA ratio = 1, and that all synaptic connections are sampled randomly with probability *p* = 0.1 (see Methods for details). Synaptic weights are constant across network sizes, with a peak EPSP of ∼ 0.6 mV at rest (NMDA/AMPA ratio, *α*_*N/A*_ = 1). The recurrent I/E-ratio, *β* = 3.1, is set by scaling all inhibitory conductances by *β* relative to the excitatory conductances (at a reference potential of *V*_0_ = −60 mV). We model the effect of NMDAR knockout in a subset of excitatory neurons, *N*_*KO*_ = 10, by eliminating their presynaptic NMDARs (NMDA/AMPA ratio = 0). The number of affected neurons is kept small to avoid potential changes in network dynamics resulting from the loss of NMDARs. Furthermore, in cells with NMDAR knockout, we assume that the remaining AMPARs have potentiated to restore the total excitatory charge of a single input (at *V*_0_ = −60 mV). Across all networks, all neurons receive independent, Poisson-distributed inputs sampled at *λ*_*ff*_ = 400 Hz. All feedforward synapses are without NMDARs.

**Figure 4.**
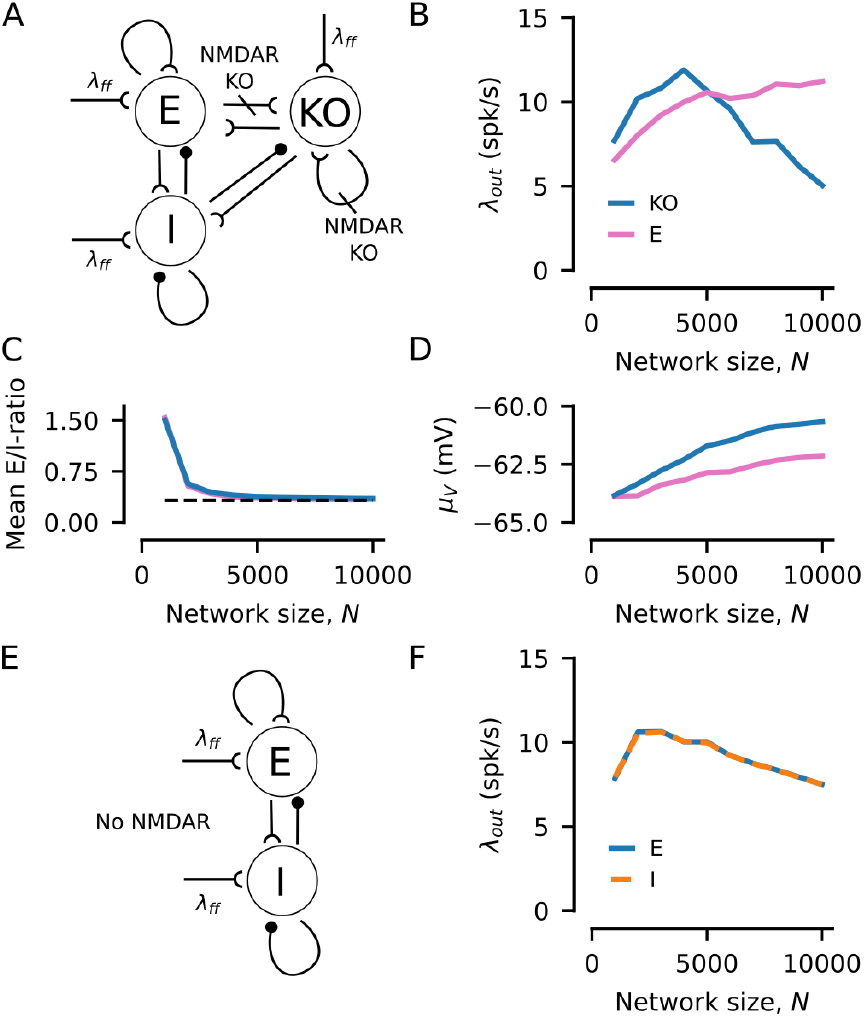
Differential effects of single-cell NMDAR knockout across the cortical hierarchy. **A**) NMDAR knockout network model. **B**) Average postsynaptic activity for each subpopulation across network sizes. **C**) Mean conductance E/I-ratio (across feedforward and recurrent synapses). Horizontal line indicates the recurrent E/I-ratio (1*/β*). **D**) Mean membrane potentials. **E**) Network model without NMDARs. **F**) Average postsynaptic activity for each subpopulation across network sizes without NMDARs. Each network was simulated for 3 s of simulated time after a 2 s warm-up period. Panels C-D: data are averaged across 10 neurons per subpopulation, removing *±* 5 ms around each spike to limit the influence of spike generations.

Comparing the postsynaptic activity of the three neuronal subpopulations suggests that eliminating NMDARs has qualitatively different effects depending on the network size (Fig. 4B; spike rasters shown in Fig. S5). When the network is small, the number of recurrent synapses is small, and the combined input to each neuron is relatively sparse. In this regime, eliminating NMDARs increases postsynaptic activity (Fig. 4B, blue). On the other hand, for larger network sizes, the number of recurrent inputs increases, leading to greater postsynaptic activity in neurons with intact NMDARs (Fig. 4B, pink). Note that the inhibitory rates closely follow the excitatory rates (E) and are therefore not shown. In NMDAR-knockout neurons, however, postsynaptic activity instead begins to decrease. These differential effects of NMDAR knockout relative to the unperturbed neurons can not be explained by differences in E/I-ratios or mean membrane potentials between spikes (Fig. 4C-D). Lastly, as a control, we observe a similar but network-wide loss of activity in large networks with only AMPARs (NMDA/AMPA ratio = 0) (Fig. 4E-F; spike rasters shown in Fig. S6).

## Discussion

The NMDA receptor has been linked to a diverse set of functions, including long-term plasticity (17; 22) and metaplasticity (42), two cellular mechanisms controlling learning and memory formation. Their voltage-dependence also has electro-physiological roles, for example, in generating dendritic action and/or plateau potentials (23; 24) and neuronal bursting, as well as functional roles (25), for example, in generating persistent activity (26). Here, we have studied the role of synaptic NMDARs on the excitability and input-output transfer function across different input levels of a thick-tuft layer 5 pyramidal cell.

We show that the increased excitability with NMDARs during depolarization influences the cell’s ability to generate action potentials in states of high presynaptic activity (Fig. 2A). Without NMDARs, postsynaptic activity not only saturates but also decreases within the tested range of inputs, similar to previously reported results (6). Thus, without NMDARs, the conductance state of a neuron limits the range of presynaptic inputs to which it can produce unique responses. However, in the presence of NMDARs, no saturation of postsynaptic activity occurs within the range of tested inputs. The lack of saturation depends on the slow increase in effective synaptic strength (e.g., Fig. S4F), and postsynaptic activity is strongly increased by the positive feedback between NMDARs and Na^+^ channels distributed across the cell (Fig. 3A-B). However, larger presynaptic inputs are needed to elicit responses. As evidence suggests an increase in both excitatory connections (7; 8; 9) and NMDAR-mediated current (21; 18; 10) across the cortical hierarchy, our results suggest that an increase in NMDAR-mediated current across the cortical hierarchy can be an important mechanism for maintaining neuronal excitability in higher cortical areas, thus providing an additional mechanism further broadening the functional repertoire of NMDARs.

A well-known consequence of high network activity is reduced input resistances (35). We show that the reduction in input resistance is significantly stronger without NMDARs (Fig. 1B). Previous studies have reported that intrinsic voltage-gated ion channels can increase the input resistance of a cell during spontaneous depolarizations (36; 37). Typically, these results rely on voltage-gated ion channels, such as inward rectifying K^+^ channels (43) or hyperpolarization-activated caution channels (44; 45), closing during depolarization. In our simulations, however, we show that when a cell receives synaptic inputs, input resistance can also increase at depolarized states by partial opening of Na^+^ channels, amplifying NMDAR-mediated currents and vice versa (Fig. 1D).

### Differential effects of NMDAR knockout/hypofunction

Our network simulations suggest that the effect of NMDAR knockout is dependent on network size (Fig. 4). Thus, we predict that NMDAR knockout will have qualitatively different effects across the cortical hierarchy. However, the results can be extended more generally to the amount of presynaptic activity, where sparse (dense) inputs potentiate (reduce) postsynaptic activity. For example, in the PFC of non-human primates, NMDAR blockade (in particular of GluN2B) reduces postsynaptic firing rates of Delay cells during oculo-motor delayed response tasks (46; 26). Furthermore, the reduction is especially strong for preferred directions. However, recordings of spontaneous activity, i.e., under low-input conditions, in rat PFC have shown that blocking NMDARs can increase postsynaptic firing rates, rather than decrease them (47). Although these observations may appear contradictory, they are consistent with our results on the differences in neuronal gain in high- and low-input regimes. Thus, our work provides a mechanism to explain the differential effects of NMDAR knockout/blockade as a function of presynaptic activity.

Furthermore, network-wide NMDAR knockout produces effects similar to those of single-cell knockouts (Fig. 4F). NMDAR hypofunction, i.e., reduced NMDAR activity, has been linked to neurological diseases such as schizophrenia (48). Thus, it is interesting to speculate about the implications of these input-dependent changes in neuronal gain, with and without NMDARs, for the neurological basis of schizophrenia. However, more detailed cortical network models (e.g., with different NMDA/AMPA ratios across neuronal subtypes) are needed to further investigate how NMDAR hypofunction affects cortical network activity.

### Systematic dendritic variations in E/I- and NMDA/AMPA-ratios

Our study suggests that including NMDARs causes stronger gain modulation with changes in the E/I ratio (Fig. 2C). This result is corroborated by a previous study showing that including NMDARs makes the neuronal gain more sensitive to increased inhibitory conductance (49). Whole-cell mappings of excitatory and inhibitory synapses in mouse cortical layer 2/3 pyramidal neurons have shown non-uniform distributions of both excitatory and inhibitory synapses, with increasing E/I ratios further from the soma (50). Thus, different parts of the dendritic tree might be differently affected by NMDARs. Furthermore, there can also be systematic differences in the NMDA/AMPA ratio across the dendritic tree. For example, layer 5 pyramidal cells in the retrosplenial cortex have high NMDA/AMPA ratios along the basal and distal apical dendrites, but not in the oblique dendrites (51). Although these findings are from cells in different layers and brain areas, they suggest the intriguing possibility that different parts of dendritic trees might be differently adapted to integrate dense or sparse inputs. In addition, non-uniform distributions of E/I- and NMDA/AMPA ratios could also increase the influence of regenerative currents (e.g., NMDA spikes).

### Regulation of NMDA/AMPA ratios

Calcium-based long-term potentiation (LTP) and depression (LTD) primarily up-or down-regulate the expression of AMPARs within excitatory synapses (17). Thus, there can be long-lasting, activity-dependent changes in the NMDA/AMPA ratio of individual synapses. Comparably little is known about the regulation of NMDARs. However, between layer 5 pyramidal cells in V1, there is a compensatory, but much slower, regulation of NMDARs during LTP (52). A similar late regulation of NMDARs has been observed in the Schaffer-CA1 synapse (53). NMDARs might also undergo changes independent of AMPAR regulation (see (54) for a review). Thus, understanding the mechanisms underlying the regulation of NMDA/AMPA ratios (whether through changes in peak conductance or changes in conveyed charge) would yield important insights into how single neurons adapt to the range of inputs they receive.

### Uncertainties in dynamics of Mg^**2+**^ block

A significant part of the postsynaptic excitability can be attributed to the interaction between the voltage-dependence of NMDARs and the voltage-gated Na^+^ channels (Fig. 3B, compare pink and gray traces) by increasing input resistance (Fig. 1B and D), thereby boosting depolarizations caused by partial openings of Na^+^ channels (Fig. 3F) and lowering the required synaptic depolarization for AP generation (Fig. 3G). The feedback interaction between NMDARs and Na^+^ channels accounts for more than 60% of the postsynaptic activity with NMDARs (Fig. S4C). However, our model assumes near instantaneous voltage-dependent Mg^2+^ blocking and unblocking (13). It has been shown that, depending on the NMDAR subunit composition, voltage-dependent Mg^2+^ unblocking also contains components with slower timescales (55; 56). Thus, our model of NMDARs might overestimate the amount of NMDAR-mediated charge conveyed for a given choice of NMDAR parameters. Furthermore, these slow components can limit NMDARs’ ability to react to fast depolarizations caused by the opening of Na^+^ channels (55), and hence limit the strength of the NMDAR-Na^+^ channel feedback. However, the kinetics of Mg^2+^ unblocking are dependent on the timing of depolarization after glutamate release, with the fast component of Mg^2+^ unblocking being more prominent in response to depolarizations early after glutamate release (57). Hence, the effect of the feedback between NMDARs and Na^+^ channels is mostly overestimated at low presynaptic rates. On the other hand, with increasing input rates, NMDAR saturation (58) could become a different limiting factor if the number of synapses (on which the inputs are distributed) is small. Importantly, NMDAR saturation might be more prominent when the decay time constant of the NMDARs is long. As both Mg^2+^ unblocking and saturation can depend on the NMDAR subunit composition, this raises the possibility of different subunit compositions being optimal for different input rates beyond the results found in Fig. 2C-D. In addition, our simulations assumed input spike trains with Poissonian statistics. However, deviations from Poissonian inputs, such as bursting, may also affect Mg^2+^ unblocking and NMDAR saturation. Lastly, there is debate over whether spines are electrically distinct, isolated compartments (e.g., 59; 60). Strong electrical isolation could further shield NMDA receptors from small depolarizations caused by the Na^+^ channels and thereby further limit feedback between them. On the other hand, strong electrical isolation also implies larger depolarizations in the spine head and, therefore, increased AMPAR efficacy in unblocking NMDARs. Thus, further investigation into how slow Mg^2+^ unblocking and NMDAR saturation are affected by different input statistics is needed to deepen our understanding of NMDAR effects across different network activity states.

### Conclusions

The amount of NMDAR-mediated charge should strongly influence the integration of information. Our work suggests that integrating information across a large range of inputs benefits from NMDARs to shift and expand the sensitive range of the neuronal gain function, without the need for other homeostatic mechanisms such as synaptic scaling or intrinsic excitability (61). Furthermore, experimental evidence suggests that a shift in NMDAR subunit composition, and thereby a shift in neuronal gain, can accompany the increase in basal spine counts along the cortical hierarchy. Therefore, our results suggest that different brain regions, or possibly even different dendritic regions, may be differently adapted to integrate dense or sparse inputs.

## Methods

### Thick-tuft layer 5 pyramidal model

The morphology and passive properties of the TTL5 model are as described in (30), but we removed the voltage-gated channels to avoid interference with calcium channels. As the reconstructed axon is incomplete, we add myelinated axonal compartments and nodes of Ranvier (2 branches with 10 myelinated compartments each), see (39) for details. We distribute 5000 excitatory (containing both AMPA and NMDA receptors) and 2500 inhibitory (GABA receptors) synapses across the dendritic tree according to the membrane surface area (Fig. S1A). In addition, we add 357 inhibitory synapses somatically (approximately 12.5% of all inhibitory synapses). However, the exact number of synapses is unlikely to affect the results if we maintain the total (i.e., summed across all synapses) excitatory and inhibitory input rates. Voltage-gated Na^+^ channels (‘Nat’) and fast, high-voltage-activated (‘Kfast’) and slow, non-inactivating (‘KM’) K^+^ channels (62) are distributed across the cell with channel densities listed in Table S1. The voltage-dependence of the axonal Na^+^ channels (‘Nax’) is adjusted based on experimental recordings as described in (39). The equilibrium potential of Na^+^ and K^+^ is set to +55 and −85 mV, respectively. Furthermore, hyperpolarization-activated cation channels (HCN) are added to the soma and dendrites. The spatial distribution of HCN conductances follows the estimated fit from (63) along the proximal dendrites, while the density at the apical tuft is capped and kept uniform (64). Synaptic time constants (explained below) are given in Table S2. These channel densities and synaptic time constants yield plausible resting-state input resistances (Fig. 1B, *λ*_*E*_ = 0 kHz) and postsynaptic firing rates (Fig. 2). The simulations are performed with the Neuron simulator (version 8.2.6) (65), at a simulation temperature of 36°C and a time step of 50 μs.

### Single-compartment model

For the network model, we use single-compartment neuron models assuming cylindrical compartments with length *L* = 25 μm and diameter *d* = 25 μm. The leak conductance is set to 0.4 mS cm^−2^ (passive *R*_*in*_ ≈ 127 MΩ), the leak reversal potential *E*_*L*_ = −70 mV, and the membrane capacitance is set such that the passive membrane time constant is 20 ms. Action potentials are generated by HH-dynamics, using Na^+^ channels (‘Nat’, 2500 pS μm^−2^) and fast, high-voltage-activated K^+^ channels (‘Kfast’, 500 pS μm^−2^). Again, we use a simulation temperature of 36°C and a time step of 50 μs. Synaptic time constants (explained below) are given in Table S2.

### Synaptic models

The synaptic conductances, *g*_*X*_ (*t*), are modeled using differences of Gaussians

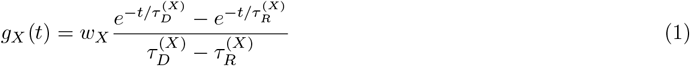

where 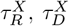,and *w*_*X*_ are the rise and decay time constants and synaptic weight (i.e., the time-integrated conductance), respectively. The time constants and reversal potentials of AMPAR, NMDAR, and GABAR are found in Table S2 (TTL5 model) and Table S3 (network model). The voltage-dependence of NMDARs is modelled as a scaling of the NMDAR-mediated conductance by a factor *m*(*V* (*t*)) using the steady-state description by (13), assuming a Mg^2+^ concentration of 1 mM,

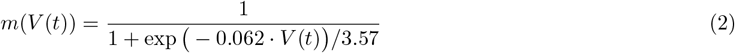

where *V* (*t*) is the local membrane potential (measured in mV) of the compartment where the synapse is situated. The NMDA/AMPA ratio, *α*_*N/A*_, is defined as the ratio of the peak conductances mediated by NMDARs and AMPARs in the absence of Mg^2+^-block

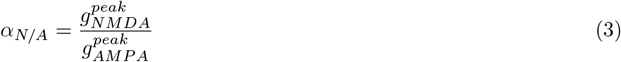

For each choice of NMDA/AMPA ratio, the synaptic weights are jointly scaled such that the combined integrated conductance (including the NMDAR’s voltage-dependence) is set to 7 nS ms measured at *V*_0_ = −54.5 mV (TTL5 model) or 2 nS ms measured at *V*_0_ = −60 mV (network model). For the TTL5 model, the integrated unitary inhibitory conductance is assumed to be a fraction 1.5 larger than the excitatory and, unless otherwise stated, the total inhibitory input rate (summed across synapses) is set such that the average E/I conductance ratio is 0.5.

## Acknowledgment

Parts of these results have been published in the doctoral thesis of M. Lenninger (66).

## Notes

### Competing Interest Statement

The authors have declared no competing interest.

### Funding

- StratNeuro

## References

[1] A. Destexhe, M. Rudolph, J. M. Fellous, and T. J. Sejnowski, “Fluctuating synaptic conductances recreate in vivo-like activity in neocortical neurons,” Neuroscience, vol. 107, pp. 13–24, 11 2001.

[2] F. S. Chance, L. F. Abbott, and A. D. Reyes, “Gain modulation from background synaptic input,” Neuron, vol. 35, pp. 773–782, 8 2002.

[3] S. A. Prescott and Y. De Koninck, “Gain control of firing rate by shunting inhibition: Roles of synaptic noise and dendritic saturation,” Proceedings of the National Academy of Sciences of the United States of America, vol. 100, pp. 2076–2081, 2 2003.

[4] S. A. Prescott, S. Ratté, Y. De Koninck, and T. J. Sejnowski, “Pyramidal neurons switch from integrators in vitro to resonators under in vivo-like conditions,” Journal of Neurophysiology, vol. 100, pp. 3030–3042, 12 2008.

[5] K. A. Ferguson and J. A. Cardin, “Mechanisms underlying gain modulation in the cortex,” Nature Reviews Neuroscience 2020 21:2, vol. 21, pp. 80–92, 1 2020.

[6] A. Kuhn, A. Aertsen, and S. Rotter, “Neuronal Integration of Synaptic Input in the Fluctuation-Driven Regime,” Journal of Neuroscience, vol. 24, pp. 2345–2356, 3 2004.

[7] B. G. Cragg and M. R. C. Cerebral, “The density of synapses and neurones in the motor and visual areas of the cerebral cortex,” Journal of Anatomy, vol. 101, p. 639, 9 1967.

[8] J. DeFelipe and I. Fariñas, “The pyramidal neuron of the cerebral cortex: Morphological and chemical characteristics of the synaptic inputs,” Progress in Neurobiology, vol. 39, pp. 563–607, 12 1992.

[9] R. Chaudhuri, K. Knoblauch, M. A. Gariel, H. Kennedy, and X. J. Wang, “A Large-Scale Circuit Mechanism for Hierarchical Dynamical Processing in the Primate Cortex,” Neuron, vol. 88, pp. 419–431, 10 2015.

[10] X. J. Wang, “Macroscopic gradients of synaptic excitation and inhibition in the neocortex,” Nature Reviews Neuroscience, vol. 21, pp. 169–178, 2 2020.

[11] M. L. Mayer, G. L. Westbrook, and P. B. Guthrie, “Voltage-dependent block by Mg2+ of NMDA responses in spinal cord neurones,” Nature1984 309:5965, vol. 309, no. 5965, pp. 261–263, 1984.

[12] L. Nowak, P. Bregestovski, P. Ascher, A. Herbet, and A. Prochiantz, “Magnesium gates glutamate-activated channels in mouse central neurones,” Nature 1984 307:5950, vol. 307, no. 5950, pp. 462–465, 1984.

[13] C. E. Jahr and C. F. Stevens, “Voltage dependence of NMDA-activated macroscopic conductances predicted by single-channel kinetics,” Journal of Neuroscience, vol. 10, pp. 3178–3182, 9 1990.

[14] G. M. Durand, Y. Kovalchuk, and A. Konnerth, “Long-term potentiation and functional synapse induction in developing hippocampus,” Nature 1996 381:6577, vol. 381, pp. 71–75, 5 1996.

[15] A. Morabito, Y. Zerlaut, B. Serraz, R. Sala, P. Paoletti, and N. Rebola, “Activity-dependent modulation of NMDA receptors by endogenous zinc shapes dendritic function in cortical neurons,” Cell Reports, vol. 38, p. 110415, 2 2022.

[16] C. I. Myme, K. Sugino, G. G. Turrigiano, and S. B. Nelson, “The NMDA-to-AMPA ratio at synapses onto layer 2/3 pyramidal neurons is conserved across prefrontal and visual cortices,” J. Neurophysiol., vol. 90, pp. 771–779, 8 2003.

[17] R. C. Malenka and R. A. Nicoll, “Long-term potentiation - A decade of progress?,” Science, vol. 285, pp. 1870–1874, 9 1999.

[18] J. B. Burt, M. Demirtaş, W. J. Eckner, N. M. Navejar, J. L. Ji, W. J. Martin, A. Bernacchia, A. Anticevic, and J. D. Murray, “Hierarchy of transcriptomic specialization across human cortex captured by structural neuroimaging topography,” Nature Neuroscience 2018 21:9, vol. 21, pp. 1251–1259, 8 2018.

[19] H. Monyer, N. Burnashev, D. J. Laurie, B. Sakmann, and P. H. Seeburg, “Developmental and regional expression in the rat brain and functional properties of four NMDA receptors,” Neuron, vol. 12, no. 3, pp. 529–540, 1994.

[20] S. Cull-Candy, S. Brickley, and M. Farrant, “NMDA receptor subunits: diversity, development and disease,” Current Opinion in Neurobiology, vol. 11, pp. 327–335, 6 2001.

[21] H. Wang, G. G. Stradtman, X. J. Wang, and W. J. Gao, “A specialized NMDA receptor function in layer 5 recurrent microcircuitry of the adult rat prefrontal cortex,” Proceedings of the National Academy of Sciences of the United States of America, vol. 105, pp. 16791–16796, 10 2008.

[22] T. V. Bliss and G. L. Collingridge, “A synaptic model of memory: long-term potentiation in the hippocampus,” Nature, vol. 361, pp. 31–39, 1 1993.

[23] J. Schiller, G. Major, H. J. Koester, and Y. Schiller, “NMDA spikes in basal dendrites of cortical pyramidal neurons,” Nature 2000 404:6775, vol. 404, pp. 285–289, 3 2000.

[24] M. E. Larkum, T. Nevian, M. Sandier, A. Polsky, and J. Schiller, “Synaptic integration in tuft dendrites of layer 5 pyramidal neurons: A new unifying principle,” Science, vol. 325, pp. 756–760, 8 2009.

[25] C. Grienberger, X. Chen, and A. Konnerth, “NMDA receptor-dependent multidendrite Ca2+ spikes required for hippocampal burst firing invivo,” Neuron, vol. 81, pp. 1274–1281, 3 2014.

[26] J. Peng, G. Wang, Y. Han, L. Qu, M. Wang, A. F. Arnsten, J. Cai, and B. Li, “Prefrontal cortical NR2B-containing NMDA receptors are essential for spatial working memory performance,” Translational Psychiatry 2025 15:1, vol. 15, pp. 1–9, 10 2025.

[27] L. M. Palmer, A. S. Shai, J. E. Reeve, H. L. Anderson, O. Paulsen, and M. E. Larkum, “NMDA spikes enhance action potential generation during sensory input,” Nature Neuroscience, vol. 17, pp. 383–390, 2 2014.

[28] S. L. Smith, I. T. Smith, T. Branco, and M. Häusser, “Dendritic spikes enhance stimulus selectivity in cortical neurons in vivo,” Nature, vol. 503, pp. 115–120, 10 2013.

[29] T. Branco, B. A. Clark, and M. Häusser, “Dendritic discrimination of temporal input sequences in cortical neurons,” Science, vol. 329, pp. 1671–1675, 9 2010.

[30] E. Hay, S. Hill, F. Schürmann, H. Markram, and I. Segev, “Models of Neocortical Layer 5b Pyramidal Cells Capturing a Wide Range of Dendritic and Perisomatic Active Properties,” PLOS Computational Biology, vol. 7, no. 7, p. e1002107, 2011.

[31] J. M. Bekkers and C. F. Stevens, “NMDA and non-NMDA receptors are co-localized at individual excitatory synapses in cultured rat hippocampus,” Nature 1989 341:6239, vol. 341, no. 6239, pp. 230–233, 1989.

[32] A. J. Watt, M. C. Van Rossum, K. M. MacLeod, S. B. Nelson, and G. G. Turrigiano, “Activity coregulates quantal AMPA and NMDA currents at neocortical synapses,” Neuron, vol. 26, pp. 659–670, 6 2000.

[33] J. J. Zhu, “Maturation of layer 5 neocortical pyramidal neurons: amplifying salient layer 1 and layer 4 inputs by Ca2+ action potentials in adult rat tuft dendrites,” The Journal of Physiology, vol. 526, pp. 571–587, 8 2000.

[34] Z. W. Zhang, “Maturation of Layer V Pyramidal Neurons in the Rat Prefrontal Cortex: Intrinsic Properties and Synaptic Function,” Journal of Neurophysiology, vol. 91, pp. 1171–1182, 3 2004.

[35] A. Destexhe, M. Rudolph, and D. Paré, “The high-conductance state of neocortical neurons in vivo,” Nature Reviews Neuroscience, vol. 4, no. 9, pp. 739–751, 2003.

[36] J. Waters and F. Helmchen, “Background Synaptic Activity Is Sparse in Neocortex,” Journal of Neuroscience, vol. 26, pp. 8267–8277, 8 2006.

[37] B. Li, B. N. Routh, D. Johnston, E. Seidemann, and N. J. Priebe, “Voltage-Gated Intrinsic Conductances Shape the Input-Output Relationship of Cortical Neurons in Behaving Primate V1,” Neuron, vol. 107, pp. 185–196, 7 2020.

[38] C. Monier, J. Fournier, and Y. Frégnac, “In vitro and in vivo measures of evoked excitatory and inhibitory conductance dynamics in sensory cortices,” Journal of Neuroscience Methods, vol. 169, pp. 323–365, 4 2008.

[39] M. H. Kole, S. U. Ilschner, B. M. Kampa, S. R. Williams, P. C. Ruben, and G. J. Stuart, “Action potential generation requires a high sodium channel density in the axon initial segment,” Nature Neuroscience 2008 11:2, vol. 11, pp. 178–186, 1 2008.

[40] M. H. Kole and G. J. Stuart, “Signal Processing in the Axon Initial Segment,” Neuron, vol. 73, pp. 235–247, 1 2012.

[41] R. Azouz and C. M. Gray, “Dynamic spike threshold reveals a mechanism for synaptic coincidence detection in cortical neurons in vivo,” Proceedings of the National Academy of Sciences of the United States of America, vol. 97, pp. 8110–8115, 7 2000.

[42] W. C. Abraham and M. F. Bear, “Metaplasticity: the plasticity of synaptic plasticity,” Trends in Neurosciences, vol. 19, pp. 126–130, 4 1996.

[43] E. S. Nisenbaum and C. J. Wilson, “Potassium currents responsible for inward and outward rectification in rat neostriatal spiny projection neurons,” Journal of Neuroscience, vol. 15, pp. 4449–4463, 6 1995.

[44] J. C. Magee, “Dendritic Ih normalizes temporal summation in hippocampal CA1 neurons,” Nature Neuroscience 1999 2:6, vol. 2, pp. 508–514, 6 1999.

[45] G. Stuart and N. Spruston, “Determinants of Voltage Attenuation in Neocortical Pyramidal Neuron Dendrites,” Journal of Neuroscience, vol. 18, pp. 3501–3510, 5 1998.

[46] M. Wang, Y. Yang, C. J. Wang, N. J. Gamo, L. E. Jin, J. A. Mazer, J. H. Morrison, X. J. Wang, and A. F. Arnsten, “NMDA Receptors Subserve Persistent Neuronal Firing during Working Memory in Dorsolateral Prefrontal Cortex,” Neuron, vol. 77, pp. 736–749, 2 2013.

[47] M. E. Jackson, H. Homayoun, and B. Moghaddam, “NMDA receptor hypofunction produces concomitant firing rate potentiation burst activity reduction in the prefrontal cortex,” Proceedings of the National Academy of Sciences of the United States of America, vol. 101, pp. 8467–8472, 6 2004.

[48] B. Moghaddam, “Bringing Order to the Glutamate Chaos in Schizophrenia,” Neuron, vol. 40, pp. 881–884, 12 2003.

[49] M. Berends, R. Maex, and E. De Schutter, “The Effect of NMDA Receptors on Gain Modulation,” Neural Computation, vol. 17, pp. 2531–2547, 12 2005.

[50] D. M. Iascone, Y. Li, U. Sümbül, M. Doron, H. Chen, V. Andreu, F. Goudy, H. Blockus, L. F. Abbott, I. Segev, H. Peng, and F. Polleux, “Whole-Neuron Synaptic Mapping Reveals Spatially Precise Excitatory/Inhibitory Balance Limiting Dendritic and Somatic Spiking,” Neuron, vol. 106, pp. 566–578, 5 2020.

[51] M. Lafourcade, M. S. H. van der Goes, D. Vardalaki, N. J. Brown, J. Voigts, D. H. Yun, M. E. Kim, T. Ku, and M. T. Harnett, “Differential dendritic integration of long-range inputs in association cortex via subcellular changes in synaptic AMPA-to-NMDA receptor ratio,” Neuron, vol. 110, pp. 1532–1546, 5 2022.

[52] A. J. Watt, P. J. Sjöström, M. Häusser, S. B. Nelson, and G. G. Turrigiano, “A proportional but slower NMDA potentiation follows AMPA potentiation in LTP,” Nature neuroscience, vol. 7, pp. 518–524, 5 2004.

[53] Y. Peng, J. Zhao, Q. H. Gu, R. Q. Chen, Z. Xu, J. Z. Yan, S. H. Wang, S. Y. Liu, Z. Chen, and W. Lu, “Distinct trafficking and expression mechanisms underlie LTP and LTD of NMDA receptor-mediated synaptic responses,” Hippocampus, vol. 20, pp. 646–658, 5 2010.

[54] D. L. Hunt and P. E. Castillo, “Synaptic plasticity of NMDA receptors: mechanisms and functional implications,” Current Opinion in Neurobiology, vol. 22, pp. 496–508, 6 2012.

[55] M. Vargas-Caballero and H. P. Robinson, “A slow fraction of Mg2+ unblock of NMDA receptors limits their contribution to spike generation in cortical pyramidal neurons,” Journal of Neurophysiology, vol. 89, pp. 2778–2783, 5 2003.

[56] R. J. Clarke and J. W. Johnson, “NMDA Receptor NR2 Subunit Dependence of the Slow Component of Magnesium Unblock,” Journal of Neuroscience, vol. 26, pp. 5825–5834, 5 2006.

[57] B. M. Kampa, J. Clements, P. Jonas, and G. J. Stuart, “Kinetics of Mg2+ unblock of NMDA receptors: implications for spike-timing dependent synaptic plasticity,” The Journal of Physiology, vol. 556, pp. 337–345, 4 2004.

[58] L. Y. Wang, “The Dynamic Range for Gain Control of NMDA Receptor-Mediated Synaptic Transmission at a Single Synapse,” Journal of Neuroscience, vol. 20, pp. RC115–RC115, 12 2000.

[59] R. Yuste, “Dendritic spines and distributed circuits,” Neuron, vol. 71, pp. 772–781, 9 2011.

[60] D. Zecevic, “Electrical properties of dendritic spines,” Biophysical Journal, vol. 122, pp. 4303–4315, 11 2023.

[61] G. G. Turrigiano, “The Self-Tuning Neuron: Synaptic Scaling of Excitatory Synapses,” Cell, vol. 135, pp. 422–435, 10 2008.

[62] Z. F. Mainen, J. Joerges, J. R. Huguenard, and T. J. Sejnowski, “A model of spike initiation in neocortical pyramidal neurons,” Neuron, vol. 15, pp. 1427–1439, 12 1995.

[63] M. H. Kole, S. Hallermann, and G. J. Stuart, “Single Ih Channels in Pyramidal Neuron Dendrites: Properties, Distribution, and Impact on Action Potential Output,” Journal of Neuroscience, vol. 26, pp. 1677–1687, 2 2006.

[64] M. T. Harnett, J. C. Magee, and S. R. Williams, “Distribution and Function of HCN Channels in the Apical Dendritic Tuft of Neocortical Pyramidal Neurons,” Journal of Neuroscience, vol. 35, pp. 1024–1037, 1 2015.

[65] N. T. Carnevale and M. L. Hines, “The NEURON Book,” The NEURON Book, pp. 1–457, 1 2006.

[66] M. Lenninger, Stimulus representation in single neurons and neuronal populations: Role of tuning shapes on minimal decoding times, and input-output functions under in-vivo-like inputs. PhD thesis, KTH Royal Institute of Technology, Stockholm, 2025.

