## Supplementary Information for "NMDAR-mediated shift of neuronal gain across the cortical hierarchy"

### Approximation of input resistance in a single-compartment model

For the single-compartment model, we model the membrane potential as

$$C\dot{V}_t = g_L(E_L - V_t) + g_{G,t}(E_G - V_t) + g_{A,t}(E_E - V_t) + f_{N,t}m(V_t)(E_E - V_t) + I_t \tag{S1}$$

where $C$ is the membrane capacitance, $E_L$, $E_G$ and $E_E$ is the leak, inhibitory, and excitatory reversal potentials, respectively, $g_L$ is the leak conductance, $g_{G,t}$ and $g_{A,t}$ are the stochastic GABAR and AMPAR conductances at time $t$, $f_{N,t}$ is the stochastic NMDAR conductance without $Mg^{2+}$-block, $m(V_t)$ is the scaling due to $Mg^{2+}$-block, and $I_t$ is the externally injected current. Note that, by Campbell's theorem, the average conductances can be approximated as

$$\begin{aligned} \mathbb{E}[g_{G,t}] &= \bar{g}_G \approx \lambda_I w_G, \\ \mathbb{E}[g_{A,t}] &= \bar{g}_A \approx \lambda_E w_A, \\ \mathbb{E}[f_{N,t}] &= \bar{f}_N \approx \lambda_E w_N \end{aligned} \tag{S2}$$

where $\lambda_E$ and $\lambda_I$ are the excitatory and inhibitory input rates, respectively, and $w_G$, $w_A$, and $w_N$ are the synaptic weights (see main text).

To approximate the input resistance, we first assume that no current is injected, i.e., $I_t = 0$. By taking the expected value of Eq. S1 we obtain the following approximation

$$0 \approx g_L(E_L - \mu_t) + \bar{g}_G(E_G - \mu_t) + \bar{g}_A(E_E - \mu_t) + \bar{f}_{N,t}\mathbb{E}[m(V_t)(E_E - V_t)] \tag{S3}$$

where $\mu_t$ is the expected value of $V_t$. Assuming the compartment is in a steady state, we have $\mu_t = \mu$. Furthermore, we approximate the expectation of the voltage-dependent $Mg^{2+}$-block by a first order Taylor expansion around $V_t = \mu$

$$\mathbb{E}[m(V_t)(E_E - V_t)] \approx m(\mu)(E_E - \mu) \tag{S4}$$

Thus, we have

$$0 \approx g_L(E_L - \mu) + \bar{g}_G(E_G - \mu) + \bar{g}_A(E_E - \mu) + \bar{f}_{N,t}m(\mu)(E_E - \mu) \tag{S5}$$

Next, we assume a steady state current injection, $I_t = I$. To simplify the calculations, we set $V_t = \delta_t + \mu$, where $\mu$ is defined as the mean membrane potential without current injection from above. Thus, $\delta_t$ corresponds to the depolarization (or hyperpolarization) caused by the current injection. This time, for the first-order approximation of the voltage-dependent $Mg^{2+}$-block around $V_t = \mu$, we have

$$\mathbb{E}[m(V_t)(E_E - V_t)] \approx m(\mu)(E_E - \mu) + m'(\mu)(E_E - \mu)\bar{\delta}_t - m(\mu)\bar{\delta}_t \tag{S6}$$

where $\bar{\delta}_t = \mathbb{E}[\delta_t]$. Thus, the expectation of Eq. S1 becomes

$$0 \approx g_L(E_L - \mu - \bar{\delta}_t) + \bar{g}_G(E_G - \mu - \bar{\delta}_t) + \bar{g}_A(E_E - \mu - \bar{\delta}_t) + \bar{f}_{N,t}\big[m(\mu)(E_E - \mu) + m'(\mu)(E_E - \mu)\bar{\delta}_t - m(\mu)\bar{\delta}_t\big] + I \tag{S7}$$

Using the approximation in Eq. S5, Eq. S7 becomes

$$0 \approx -g_L\bar{\delta}_t - \bar{g}_G\bar{\delta}_t - \bar{g}_A\bar{\delta}_t + \bar{f}_{N,t}\big[m'(\mu)(E_E - \mu)\bar{\delta}_t - m(\mu)\bar{\delta}_t\big] + I \tag{S8}$$

By solving for $\bar{\delta}_t$, assuming a steady-state, we get

$$\bar{\delta}_t = \bar{\delta} = \frac{I}{g_L + \bar{g}_G + \bar{g}_A + \bar{f}_N\big(m(\mu) - m'(\mu)(E_E - \mu)\big)} \tag{S9}$$

and thus, the approximated input resistance becomes

$$R_{in} = \frac{\bar{\delta}}{I} = \frac{1}{g_L + \bar{g}_G + \bar{g}_A + \bar{f}_N\big(m(\mu) - m'(\mu)(E_E - \mu)\big)} \tag{S10}$$

Eq. S10 shows two important results. First, for a small injected current, the input resistance is independent of the current sign (depolarizing/hyperpolarizing), regardless of the NMDA/AMPA ratio. Second, NMDARs are expected to increase the input resistance by altering the synaptic conductances proportional to $m'(\mu)(E_E - \mu)$. We verify the sign-independence in Fig. S2.

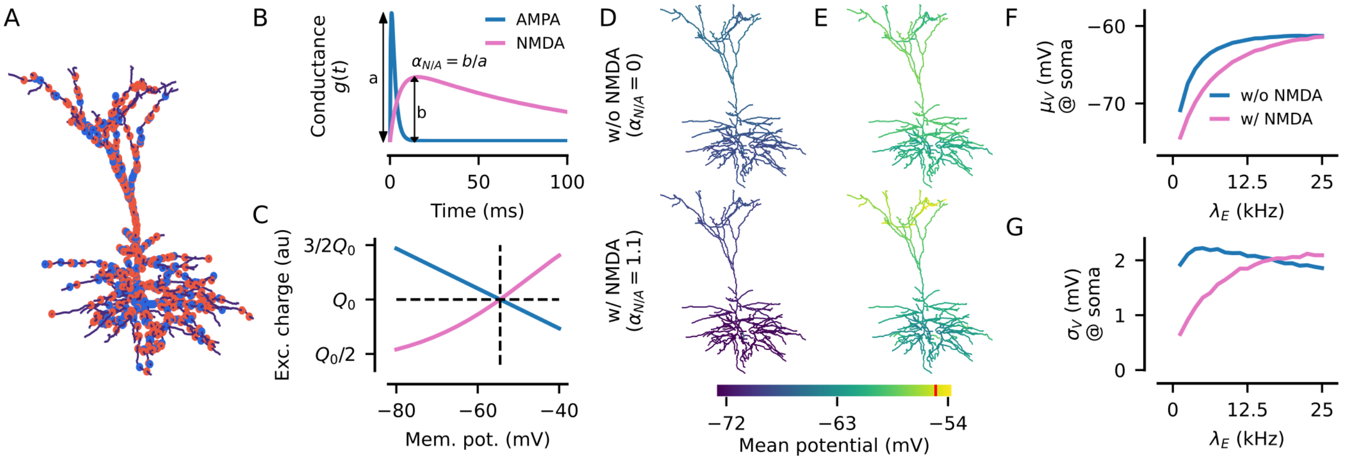


Figure S1: Properties of AMPA and NMDA receptors. **A**) Simulated TTL5 cell with excitatory (red dots) and inhibitory (blue dots) synapses distributed across the cell (10% of all synapses shown). **B**) Kinetics of AMPA (blue) and NMDA (pink) receptors. The NMDA/AMPA ratio, $\alpha_{N/A}$, is the ratio of respective peak conductances. **C**) Differences in voltage-dependence of the effective synaptic weight (measured as total charge) for NMDA/AMPA ratios $\alpha_{N/A} = 0$ (blue) and $\alpha_{N/A} = 1.1$ (pink). **D-E**) Mean membrane potentials across the dendritic tree (in the absence of $Na^+$ channels) at $\lambda_E = 2$ kHz (panel D) and $\lambda_E = 10$ kHz (panel E). Red mark indicates $V_0 = -54.5$ mV. **F-G**) Mean (panel F) and standard deviation (panel G) of the subthreshold somatic membrane potential (measured without $Na^+$ channels). Each data point is estimated based on 60 seconds of simulated time.

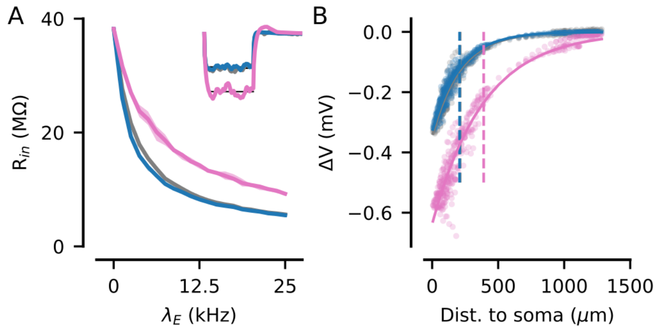


Figure S2: Input resistance with and without NMDARs is independent of current sign. **A**) Estimated input resistance versus synaptic input rate using hyperpolarizing current pulses (40 pA for 100 ms, all cells are without $Na^+$ channels). **B**) Compartment hyperpolarizations due to a single current pulse ($\lambda_E = 12.5$ kHz). Solid lines show exponential fits, and dashed lines show the characteristic lengths.

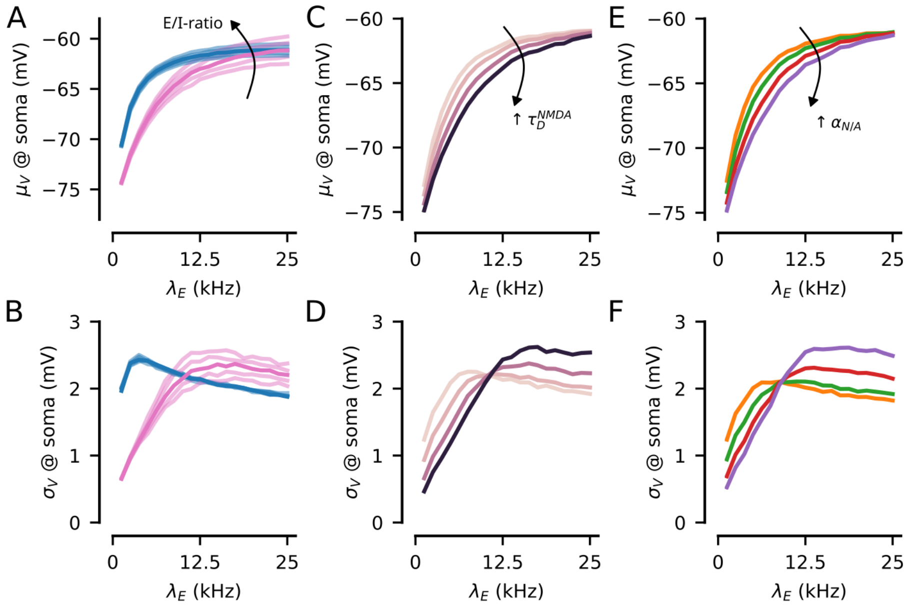


Figure S3: Subthreshold statistics for different E/I- or NMDA/AMPA-ratios. **A**) Estimated average somatic membrane potentials for different E/I ratios (explained in the main text). **B**) Corresponding estimates of the standard deviations of the somatic membrane potentials. **C - D**) Same as panels A)-B) but for various NMDA conductance decay times (explained in the main text). **E - F**) Same as panels A)-B) but for various peak NMDA/AMPA ratios (explained in the main text). All panels: data estimated from voltage recordings in the soma with $\pm 5$ ms removed around each spike.

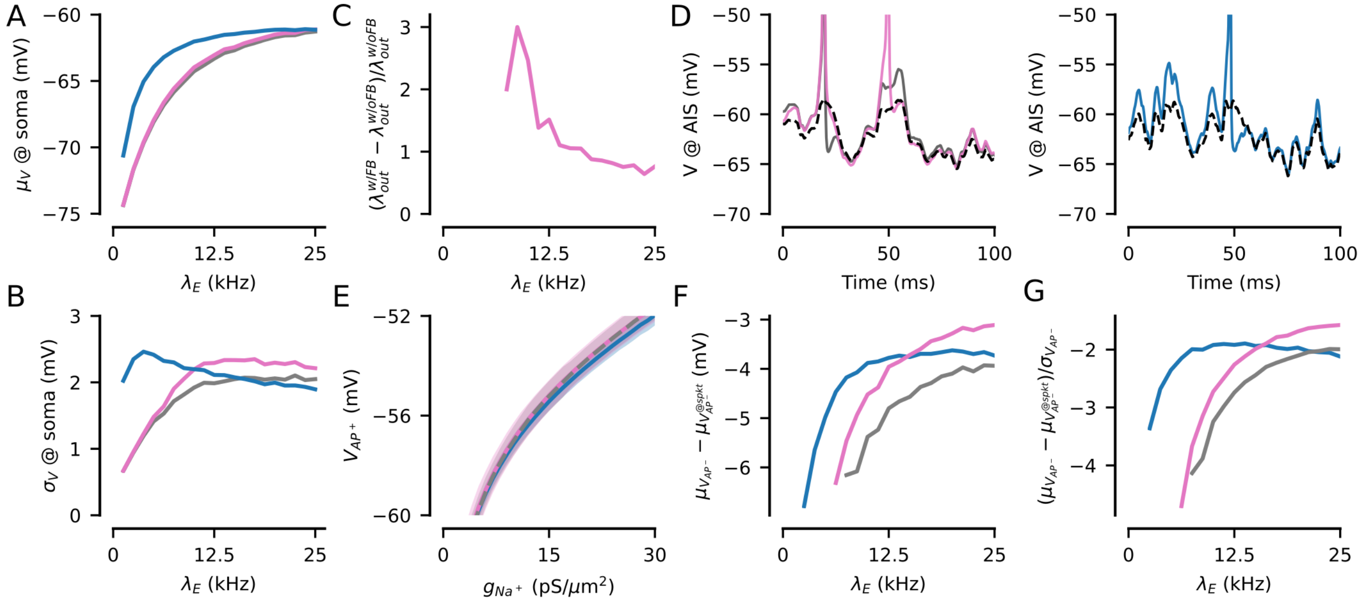


Figure S4: NMDAR - $Na^+$ channel feedback increases postsynaptic activity - Supplementary figures. **A**) Average somatic membrane potentials at the soma in $AP^+$ condition, voltage data removed $\pm 5$ ms around each spike. Blue - w/o NMDA, pink - w/ NMDA, and gray - w/ NMDA but w/o feedback. **B**) Same as panel A), but for the standard deviations of the somatic membrane potentials. **C**) Gain of including NMDA - $Na^+$ channel feedback (w/ FB) relative to the simulations where the feedback loop has been interrupted (w/o FB). **D**) (left) Voltage traces in the AIS with synaptic NMDA. Solid lines: with $Na^+$ channels ($AP^+$ condition, pink line - w/ FB, gray line - w/o FB), dashed lines: without $Na^+$ channels ($AP^-$ condition). (right) Same as the left panel but without NMDA. **E**) Mean and standard deviations of the voltage potentials (in the AIS) in $AP^+$ conditions versus the conductance of $Na^+$ channels in the AIS. Here, the gray line is dashed only to make the overlaying traces visible. **F**) Distance between the mean subthreshold potentials and the mean required synaptic depolarizations for AP generation (in the AIS). **G**) Relative distance between mean subthreshold potentials and the mean required synaptic depolarizations for AP generation, divided by the standard deviation of the subthreshold membrane potentials.

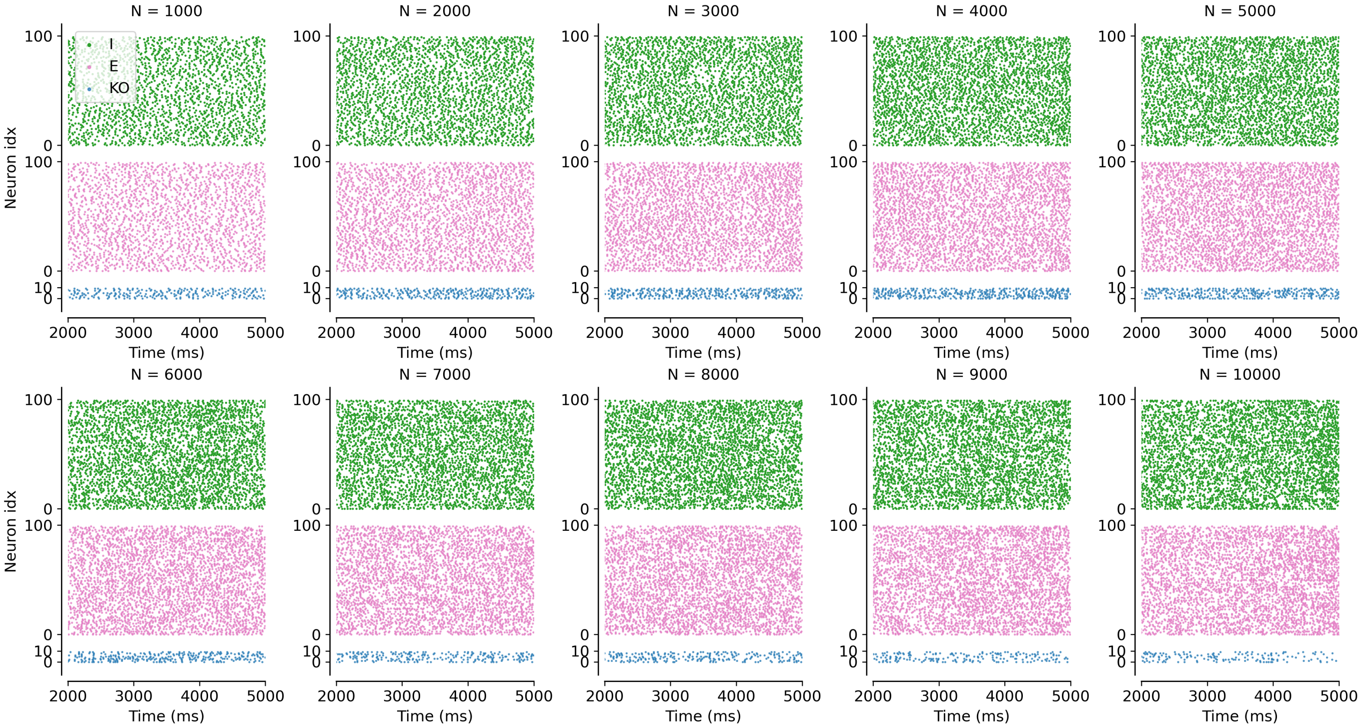


Figure S5: Spike rasters for each network size with NMDARs and NMDAR knockout (KO).

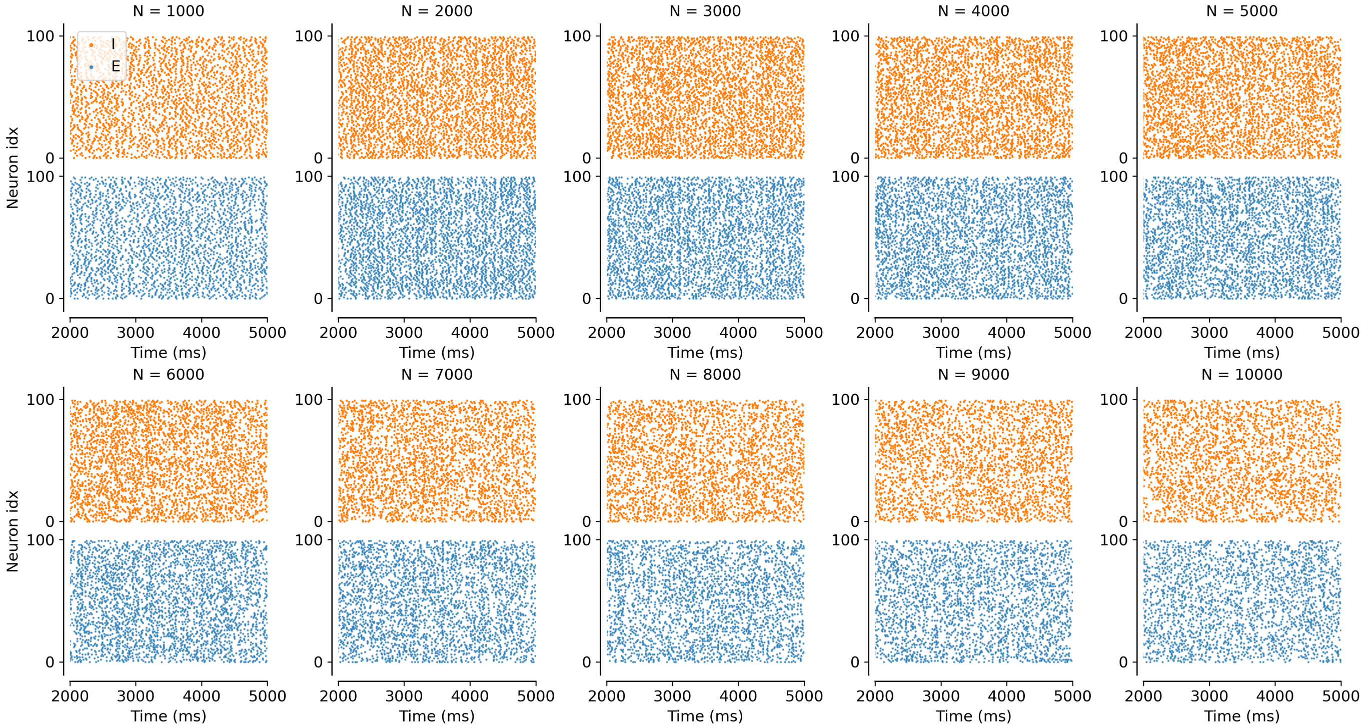


Figure S6: Spike rasters for each network size without NMDARs.

| | Soma | AIS | Dend | Nodes of Ravier | Myelinated seg. |
|---|---|---|---|---|---|
| Nat (pS µm$^{-2}$) | 100 | N/A | 45 | N/A | N/A |
| Nax (pS µm$^{-2}$) | N/A | 2500 | N/A | 2500 | 85 |
| $K_{fast}$(pS µm$^{-2}$) | 40 | 2000 | 40 | 2000 | 40 |
| $K_M$ (pS µm$^{-2}$) | 5 | 50 | 5 | 50 | 5 |

Table S1: Voltage-gated ion channel densities of the TTL5 cell model

| | $\tau_R$ (ms) | $\tau_D$ (ms) | $E_{rev}$ (mV) |
|---|---|---|---|
| AMPA | 0.5 | 2 | 0 |
| NMDA | 5 | 100 | 0 |
| GABA | 0.5 | 6 | -80 |

Table S2: Synaptic parameters of the TTL5 cell model

| | $\tau_R$ (ms) | $\tau_D$ (ms) | $E_{rev}$ (mV) |
|---|---|---|---|
| AMPA | 0.5 | 4 | 0 |
| NMDA | 5 | 100 | 0 |
| GABA | 0.5 | 4 | -80 |

Table S3: Synaptic parameters in network model
